# Genetically Programmable Adhesive Protein Hydrogels for Sealing Perforating Corneal Trauma

**DOI:** 10.1101/2025.05.26.656098

**Authors:** Ling Xu, Ping Hu, Yaling Yu, Yutong Huang, Yuhong Ye, Hao Chen, Hong Kiu Francis Fok, Pan Zhao, Xue Wu, Songzi Kou, Zhengbing Zhou, Fei Sun, Da Long

## Abstract

The cornea, situated at the forefront of the eyeball, is often subjected to varying degrees of mechanical laceration injuries due to various traumatic incidents. Timely and precise suturing of corneal wounds to prevent the leakage of intraocular contents is a significant clinical challenge. Although some studies have attempted to use hydrogels to accurately cover corneal injuries, most of these hydrogels are only suitable for adhering to focal stromal defects that do not involve the corneal stroma. Here, a fully protein-based hydrogel, based on SpyTag-SpyCatcher chemistry, was investigated for repairing corneal perforating injury models in rats and rabbits. The hydrogel formation relies on covalent bonding between SpyTag (A) and SpyCatcher (B) under physiologically mild conditions, eliminating requirements for organic solvents, elevated temperatures, or photoactivation Notably, integration of recombinant mussel foot protein 3 (Mfp3) endowed the hydrogels with exceptional water-resistant adhesiveness, while maintaining their original mechanical properties and biocompatibility. This study demonstrates the great potential of Mfp hydrogels in the timely and precise repair of corneal perforating injuries in complex physicochemical environments.

**Teaser:** Bioinspired protein hydrogel rapidly seals corneal perforations, minimizing scarring and inflammation without sutures in animal models.

## Introduction

Open globe injuries are one of the significant causes of vision loss worldwide, with corneal injuries accounting for approximately 4.06% of blindness across all age groups^[1]^. Among these, corneal perforating injuries constitute 33.3% of all corneal injuries^[2]^. As a type of perforating injury^[3]^, corneal perforating injuries leave the eyeball exposed, making it susceptible to infections and exacerbating damage, often leading to traumatic cataracts^[4]^, sympathetic ophthalmia ^[5]^, and vision loss^[6]^. Therefore, timely closure of corneal lacerations^[7]^ is fundamental to promoting rapid corneal healing and preventing subsequent complications.

Currently, the treatment for corneal perforating injuries is limited to surgical suturing, with the success of suturing the wound edges heavily dependent on the surgeon’s skill. Traditional corneal suturing involves pulling the tissue on both sides of the wound toward the center to achieve closure. However, this process can cause corneal deformation, manifesting as flattening at the suture site, displacement of the corneal apex away from the suture, steepening of the corneal center, and irregular corneal astigmatism^[8]^. The sutures, acting as foreign bodies, may induce inflammatory reactions that affect the corneal endothelial pump function, leading to persistent corneal edema^[9]^, impaired wound healing, corneal neovascularization, and ultimately, corneal scarring. With the emergence of hydrogels as a novel material, their injectability, in situ gelation, minimally invasive nature, and tunability have made them increasingly popular for repairing damaged corneas.

Cyanoacrylate glue, with its optimal adhesive strength and polymerization rate, was initially used as a corneal patch. However, due to its non-degradability and potential to induce inflammatory reactions, corneal neovascularization, foreign body reactions, and tissue necrosis, it has gradually been replaced by fibrin sealants^[10]^. Fibrin sealant (Tisseel, Baxter, Deerfield, IL, USA), approved by the FDA as a hemostatic adjunct in cardiac surgery^[11]^, was found to be unable to adhere solely to the corneal surface in wound adhesion tests^[12]^. Recently, a polyethylene glycol (PEG) hydrogel named ReSure (Ocular Therapeutix, Inc.) was approved for intraoperative management of clear corneal incision (CCI) cataract surgeries up to 3.5 mm in length^[13]^. However, this product had its PMA certificate withdrawn in 2018 due to high rates of endophthalmitis and significant ocular adverse events.

Mussel foot proteins (Mfps), particularly Mfp3 from *Mytilus edulis*, are characterized by their remarkable underwater adhesion due to catechol-rich motifs, making them highly suitable for biomedical applications like corneal repair where robust adhesion in hydrated environments is essential^[14–16]^. To harness this capability within a controlled and adaptable system, we utilize SpyTag/SpyCatcher chemistry, a protein ligation technology that enables rapid, site-specific covalent crosslinking under physiological conditions^[16,17]^. This system allows the creation of fully protein-based hydrogels with modular architectures^[15]^.

By integrating Mfp3 into these Spy network hydrogels, we develop genetically programmable adhesives that combine Mfp3’s water-resistant adhesion with the structural precision of Spy networks. These hydrogels exhibit tunable mechanical properties, strong interfacial adhesion, and biocompatibility^[18]^, addressing the dual demands of rapid wound sealing and tissue regeneration in dynamic corneal environments. This study underscores the potential of genetically engineered protein hydrogels to transcend the limitations of conventional sutures, offering a versatile platform for precision repair of corneal perforations and advancing the development of adaptive biomaterials for ocular trauma and broader biomedical applications.

## Results

### Synthesis of Adhesive Protein Hydrogels with Tunable Mechanical Properties

This study biosynthesized the recombinant AAA and BMB proteins using *E. coli* expression systems, purified via Ni-NTA affinity chromatography, and subsequently dialyzed the proteins against Milli-Q water prior to lyophilization ^[14,15]^. We cloned and produced the copper(II)-dependent oxidative enzyme, *S. antibioticus* tyrosinase, in recombinant form using *E. coli* expression systems, followed by *in vitro* reconstitution to mimic its native catalytic activity toward tyrosine residues. We prepared AAA and BMB protein solutions at equal weight concentrations (8% and 10% in PBS), mixed them in a 1:4 volume ratio, and added tyrosinase to the mixture at a concentration of 1 μg per 30 μL. Rheological analysis revealed that both hydrogels formed within approximately 1500 seconds, demonstrating rapid gelation kinetics (Fig. 2A). We further conducted dynamic rheological tests in frequency and strain sweep modes (Fig. 2B and 2C), which showed that both hydrogels exhibited a stable storage modulus *G’* and a significantly lower loss modulus *G’’* across the shear frequency range of 0.1–100 rad/s, confirming the establishment of covalently cross-linked polymer networks. Notably, the 10 wt% hydrogel (10AB) displayed superior mechanical properties compared to the 8 wt% hydrogel (8AB), attributable to its higher protein concentration, indicating that hydrogel stiffness can be precisely modulated by adjusting protein content.

**Fig. 1.**
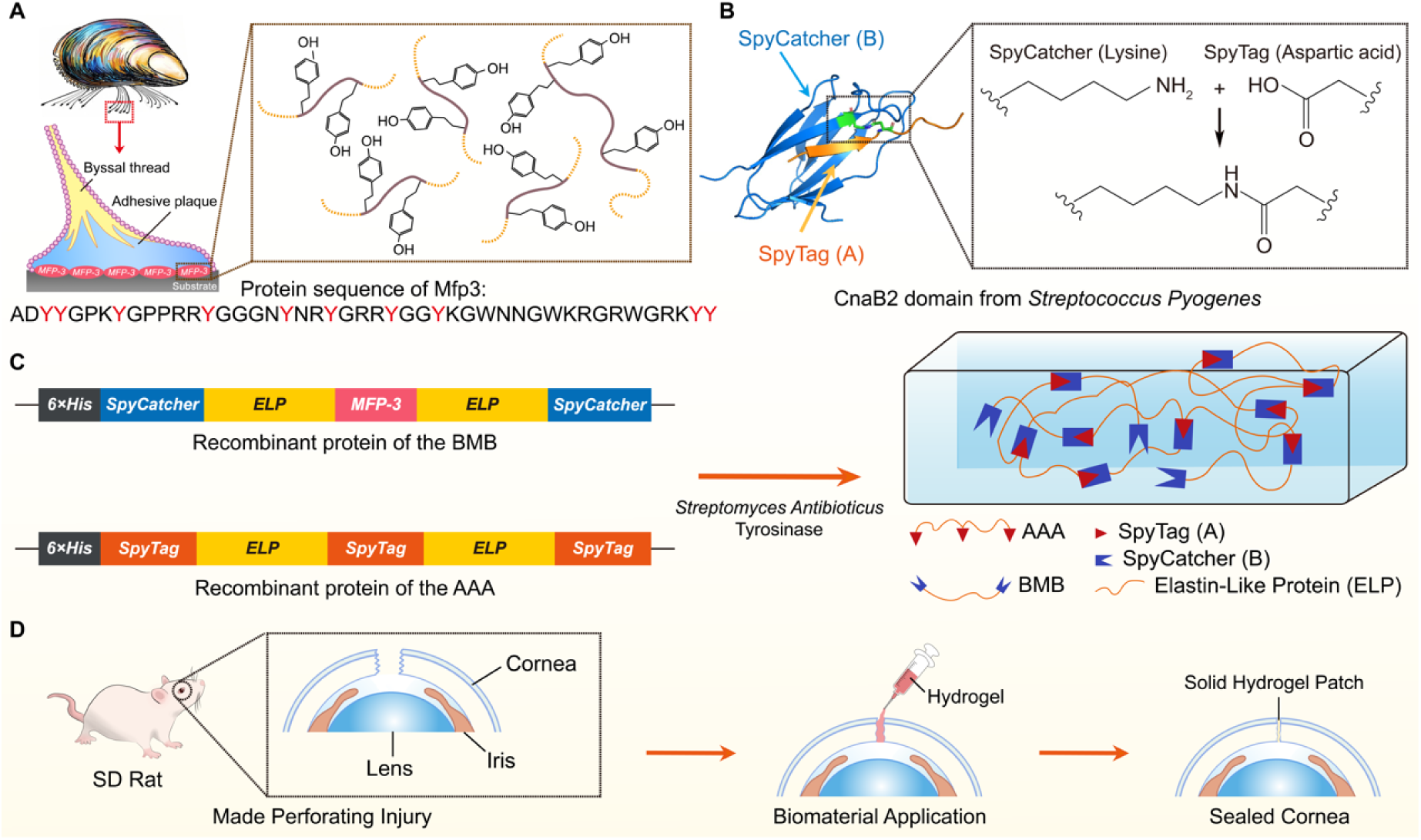
Synthesis of adhesive protein hydrogels via SpyTag-SpyCatcher chemistry and enzymatic oxidation and their application in repairing corneal perforating injuries. **(A)** Diagram of a mussel attached to a substrate, displaying the byssal thread and adhesive plaque. Alongside, the amino acid sequence of Mfp3 is shown, with tyrosine residues highlighted in red. **(B)** Structure of the SpyTag/SpyCatcher complex and the formation of isopeptide bond [Protein Data Bank identifier (PDB ID): 4MLI]. **(C)** Constructs of the SpyTag-ELP-SpyTag-ELP-SpyTag (AAA) and SpyCatcher-ELP-Mfp3-ELP-SpyCatcher (BMB) proteins composing the adhesive protein hydrogels (denoted as “Mfp hydrogels”), along with their self-assembly and subsequent gelation behavior following enzymatic oxidation. **(D)** Schematic illustration of the procedure for generating a corneal perforating injury model in a rat eyeball and sealing the wound using Mfp hydrogel adhesives.

**Fig. 2.**
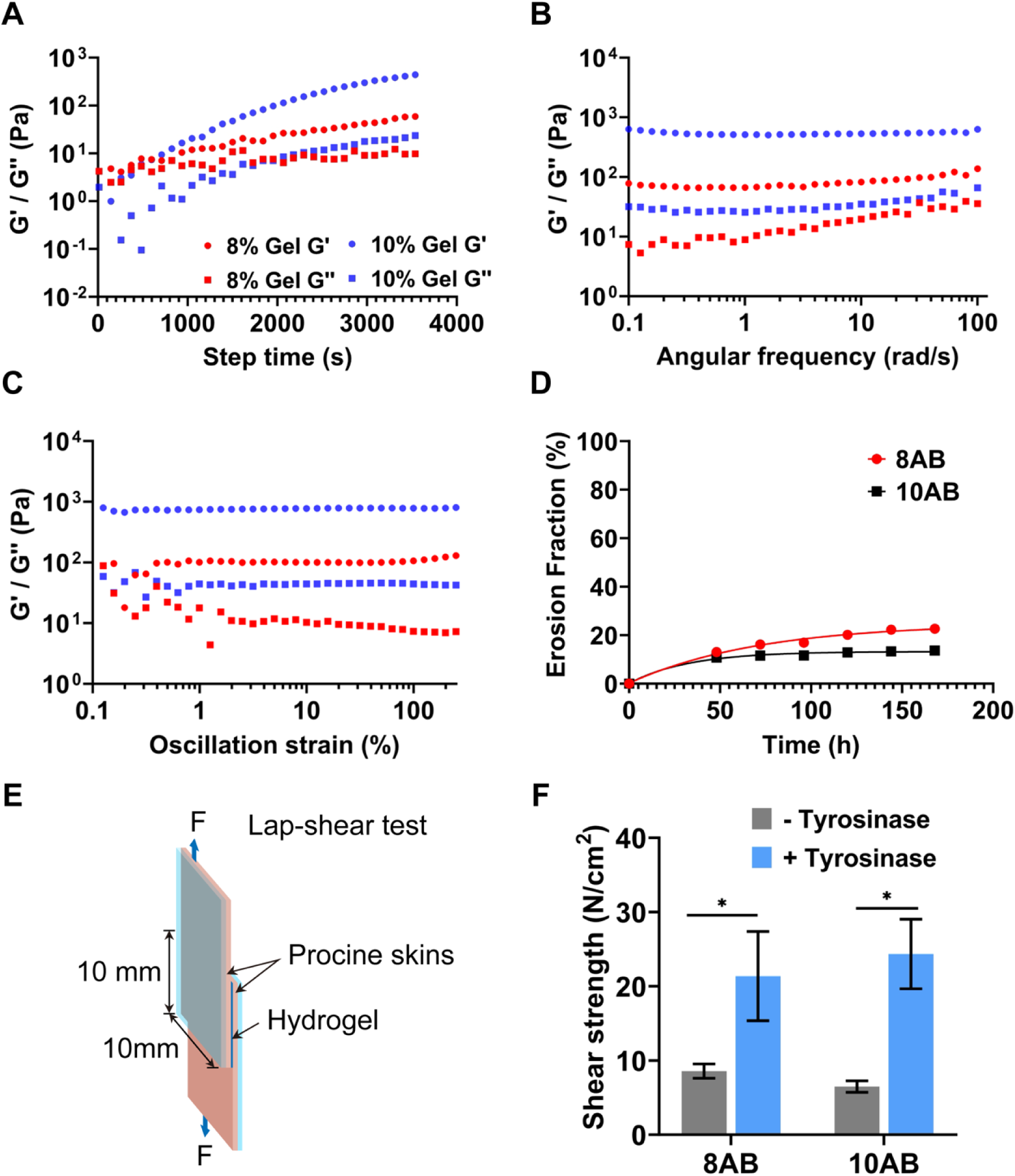
Mechanical and adhesive properties of Mfp hydrogels. **(A)** Dynamic time sweep rheology of 8AB and 10AB hydrogels at 5 rad/s and 5.0% strain (G′: storage modulus; G′′: loss modulus). 8AB and 10AB denote hydrogels composed of AAA + BMB at 8 wt% and 10 wt%, respectively (molar ratio 2:3). **(B)** Dynamic frequency sweep tests of 8AB and 10AB. **(C)** Dynamic strain sweep tests of 8AB and 10AB. **(D)** Erosion profiles of 8AB and 10AB in DPBS (pH 7.4, 37°C) over 180 h (n = 3). The data were polynomial fitted. **(E)** Adhesive strength test setup: the Mfp hydrogel bonds two tissue slides (10 mm × 10 mm contact area). **(F)** Interfacial adhesive strength quantified via tensile testing, demonstrating strong adhesion to porcine skin. A tyrosinase-free AAA/BMB mixture served as the control. Data: mean ± SD (n = 3); two-sample *t*-test, *P* < 0.05.

To evaluate *in vitro* degradation, we immersed the hydrogels in DPBS (pH 7.4) at 37°C (Fig. 2D). Erosion tests demonstrated that 10AB exhibited less than 15% mass loss after 7 days, highlighting its exceptional stability, whereas 8AB showed approximately 22% erosion due to its lower cross-linking density.

Using shear strength adhesive testing, we further assessed the adhesiveness of these protein hydrogels (Fig. 2E). With the presence of tyrosinase, both the 10AB and 8 AB hydrogels exhibited significant adhesiveness on porcine skin, which can generate adhesion strengths of 24.3±4.7 and 21.4±6.0 N/cm^2^ respectively, substantially higher than those in the absence of tyrosinase (6.5±0.8 and 8.6±1.0 N/cm^2^), suggesting the hydrogels can serve as an effective adhesive for biomaterials (Fig. 2F).

### Influence of Mfp hydrogels on limbal stem cells (LSCs)

Limbal stem cells (LSCs) line was applied to explore the cytocompatibility of the hydrogels. This study employed a 3D on-top culture model to cultivate LSCs (Fig. 3A), aiming to simulate the *in vivo* state where LSCs are activated upon injury and subsequently proliferate/migrate through hydrogels^[19]^. LSCs cultured on 8AB and 10AB hydrogels exhibited similar cell viability at all time points (1-, 3-, 5-, and 7-days post-culture) (Fig. 3B). Even on day 7, the cell survival rates on the surfaces of 8AB (∼99.54±0.33%) and 10AB (∼99.61±0.28%) hydrogels remained high (Fig. 3C). Compared to the control group (∼99.62±0.33%), the results indicated that the gel polymers formed by tyrosinase-oxidized BMB and AAA had no significant cytotoxic effect on LSCs. Additionally, although there was no statistical difference in cell viability between the two hydrogel groups with different concentrations, the overall viability of 8AB was slightly lower than that of 10AB. The CCK-8 assay results were consistent with the Calcium-AM/PI staining (Fig. 3D). Though no significant statistical difference between the 8AB hydrogel and 10AB hydrogel, the 10AB hydrogel-cultured LSCs exhibited faster multiplication rate at each time point. These data demonstrated that Mfp hydrogels possessed excellent cell compatibility and higher concentrations of AAA and BMB correlated with increased LSC activity.

**Fig. 3.**
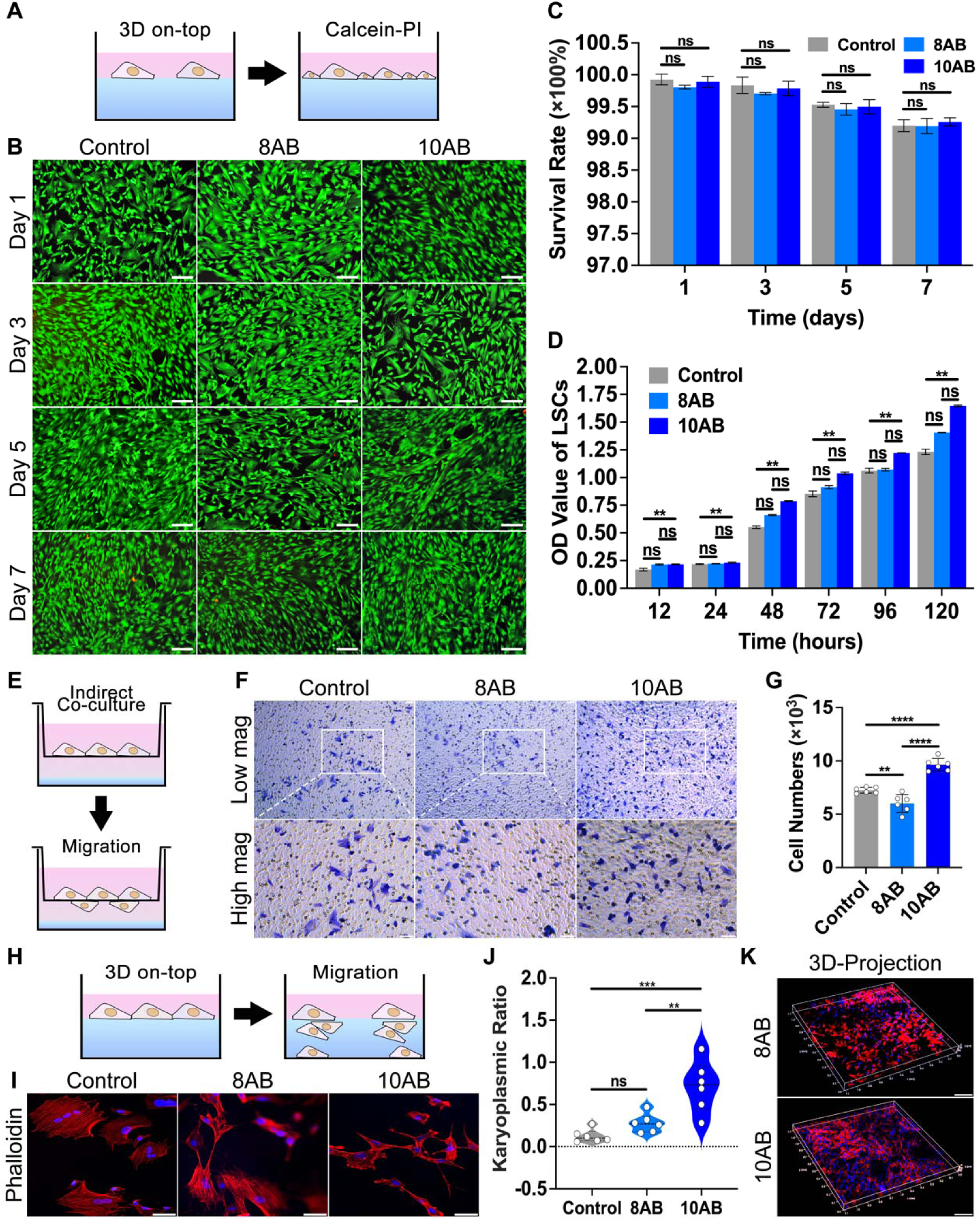
Influence of Mfp hydrogels on limbal stem cells (LSCs) **(A)** Schematic illustration of the 3D on-top culture system in live/dead staining for LSCs. The Mfp hydrogels are represented by blue rectangles, and the medium is indicated by the pink ones. **(B)** Fluorescence micrographs of LSC viability on hydrogel surfaces stained with calcein-AM (live cells, green) and propidium iodide (dead cells, red) over 7 days. Scale bar: 100 µm. **(C)** Quantification of LSC viability on 8AB, 10AB, and control surfaces from live/dead assays (n = 6). **(D)** CCK-8 assay demonstrating LSC proliferation on 8AB and 10AB over 120 hours compared to K-SFM control (n = 3). **(E)** Schematic illustration of the indirect co-culture system in Transwell assays for LSCs. **(F)** Representative images of LSC migration in Transwell assays after 24-hour culturing with PBS, 8AB, or 10AB. Upper panel: low magnification; lower panel: high magnification. Scale bar: 50 µm. **(G)** Quantification of LSC migration through Transwell membranes (n = 6). **(H)** Schematic illustration of the 3D on-top culture system in migration assays for LSCs. **(I)** Cytoskeletal organization of LSCs in 3D on-top culture visualized by phalloidin staining. Scale bar: 50 µm. **(J)** Quantitative analysis of nucleus-to-cytoplasm (N/C) ratios from phalloidin-stained samples (n = 6). Statistical analysis by two-way ANOVA; data presented as mean ± SD. Significance levels: ns = not significant, \*\**P* < 0.01, \*\*\**P* < 0.001, \*\*\*\**P* < 0.0001. **(K)** Confocal 3D reconstructions of LSCs migrating within 1 mm-thick 8AB and 10AB hydrogels. Scale bars: 150 µm (left), 200 µm (right).

Cell migration is essential for corneal repair and regeneration. Vast extracellular signals, including biochemical factors (haptotaxis or chemotaxis) and biophysical cues (topotaxis or durotaxis) can guide cellular migration^[20]^. Herein, indirect co-culture migration analysis was applied to explore the chemotaxis of Mfp hydrogels on LSCs (Fig. 3E). Compared with the blank control (∼7.248±0.28×10^3^), the LSCs treated with 10AB showed more numbers of migration cells (∼9.634±0.6×10^3^) (Fig. 3F). However, the 8AB group (∼6.013±0.84×10^3^) showed fewer numbers of vertical migrated cells than the control group (Fig. 3G). The evidence indicated that the 8AB hydrogel, exhibiting a low storage modulus (∼100 Pa), demonstrated insufficient rigidity, thereby constraining traction forces and diminishing cell migration relative to the control group. While the 10AB hydrogel with a higher storage modulus (∼1000 Pa), exhibited enhanced stiffness, facilitating increased cellular forces and providing a greater abundance of adhesion sites and growth factors, which collectively augmented cell migration^[21–23]^. To further clarify the impact of 8AB and 10AB on cell migration, a phalloidin staining experiment was conducted, using the nucleus-to-cytoplasm (N/C) ratio as a quantitative indicator (Fig. 3H and 3I). The results showed that the N/C ratios for both 8AB (0.278±0.11) and 10AB (0.714±0.3) were higher than that of the control group (0.125±0.08). A higher N/C ratio indicates stronger stem cell activity^[24]^, and the cell morphology observed under fluorescence microscopy confirmed that the gels had no adverse effects on maintaining cell morphology (Fig. 3J). High-magnification confocal microscopy 3D reconstructions (40×) were used to observe cell migration within the gels, revealing that the migration distances between the 8AB and 10AB groups were similar (Fig. 3K). Since both 8AB and 10AB have their respective advantages and exhibit no cytotoxicity, both were selected as candidate materials for subsequent *in vivo* animal experiments.

### Adhesion and Wound Healing in Perforating Corneal Trauma of Sprague-Dawley Rats

Before using New Zealand white rabbits as the experimental model, we first conducted preliminary efficacy validation on Sprague-Dawley (SD) rats to assess the usability, safety, and initial therapeutic effects of Mfp hydrogels. A total of 225 rats were included in a 2-month *in vivo* follow-up study to compare wound healing outcomes among the no-intervention (control group), suture group, and hydrogel wound closure group. Since patients typically require 3–6 months of healing time after corneal trauma suturing to meet suture removal criteria, and complete corneal healing takes 6 months or longer, these rats were further monitored for an additional 4 months (6-month endpoint) to more carefully examine the long-term effects on wound healing mediated by the Mfp hydrogels.

For each surgical model, an ophthalmic measuring gauge was used to control the corneal wound diameter at 3 ± 0.5 mm to prevent damage to the limbal LSCs region^[25]^, which could affect corneal epithelial regeneration. Subsequently, a full-thickness laceration was created using microforceps to simulate the most common clinical presentation of corneal perforating injuries (Fig. 4A). In the hydrogel group, after inducing the wound, the BMB and AAA solutions were mixed in a 1.5 mL Eppendorf tube for 5 minutes to allow the mixture to be fully oxidized by tyrosinase into an amber-hued transparent gel state. The gel mixture was then applied to the wound using ophthalmic microforceps, aligning the corneal wound edges until the hydrogel completely solidified. Adhesion efficacy was validated through absence of postoperative aqueous humor leakage, confirming stable interfacial integration.

**Fig. 4.**
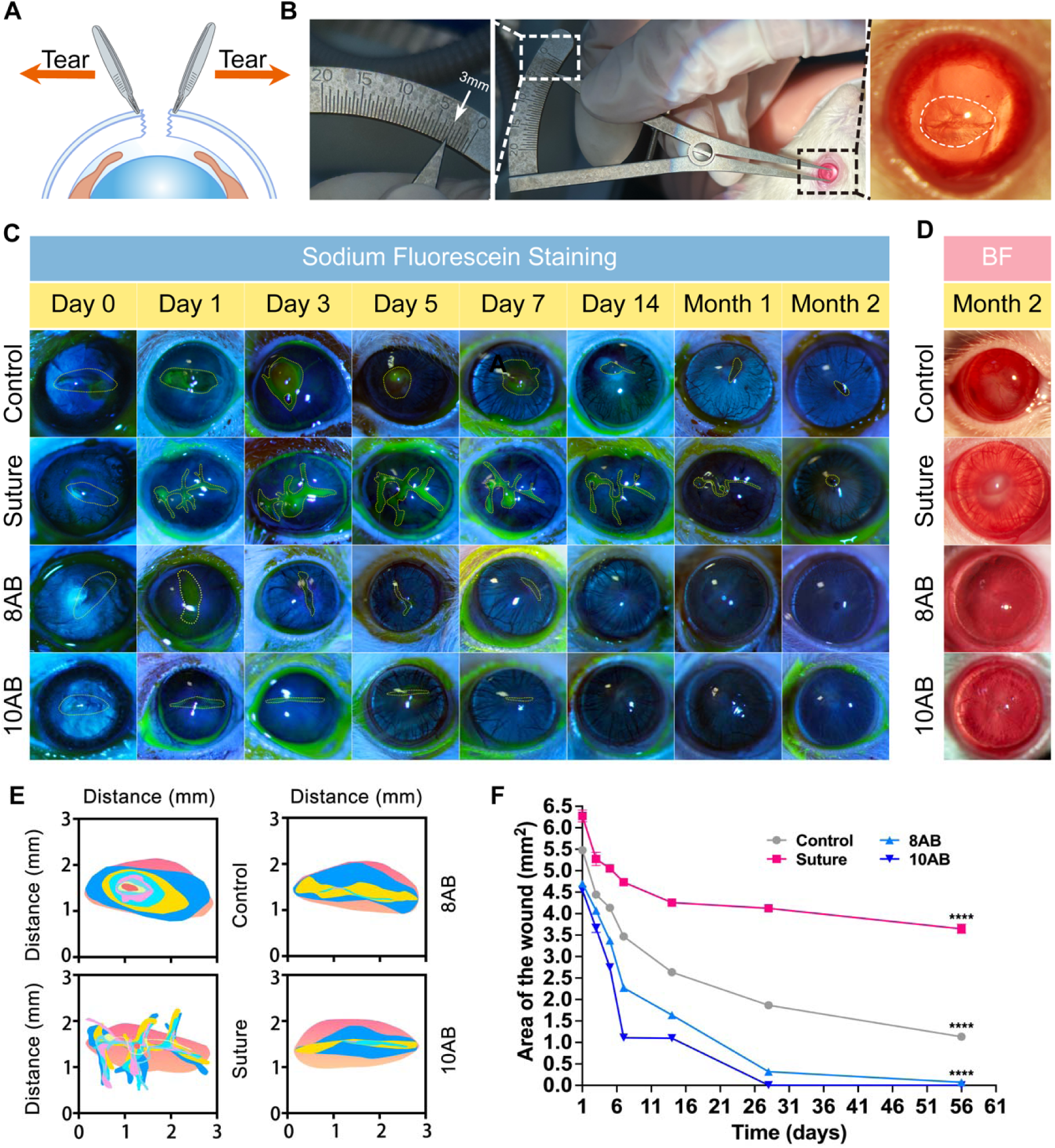
Mfp hydrogels promote wound healing in a rat corneal perforation injury model. **(A)** Schematic illustration of the procedure for generating a corneal perforating injury model in a rat eyeball using microscopic tweezers. **(B)** Corneal perforation surgery (middle), wound diameter measurement using an ophthalmic gauge (left), and representative anterior segment images immediately post-injury (right; wound margins marked by white dashed lines). **(C)** Serial fluorescein sodium staining under cobalt blue light showing wound healing in control, suture, 8AB, and 10AB groups (yellow dashed lines indicate wound borders). **(D)** Anterior segment photographs at 2 months post-operation under ambient white light. BF=bright field. **(E)** Traces of corneal wound closure during the observed 2 months. **(F)** Quantitative analysis of corneal wound area reduction, measured by fluorescein sodium staining (n=6). Statistical significance versus control group was determined by two-way ANOVA (\*\*\*\**P*<0.0001).

During the follow-up period, the corneas of the model group rats showed gradual iris prolapse 5 days post-surgery. Due to the organization of exudates and corneal repair processes, the prolapsed iris became permanently fixed within the cornea, forming an adherent leukoma to seal the corneal defect. We visually assessed the corneal surfaces of the 8AB and 10AB groups at 1-, 3-, 5-, and 7-days post-surgery for signs of corneal neovascularization and leukoma, which might indicate adverse inflammatory reactions to the hydrogel material. Although some acute inflammatory signs appeared within the first few hours (less than 24 hours) post-surgery, the irritation quickly subsided (Fig. S1). Evaluations at 3 days post-surgery showed no congestion or inflammation at the wound site, and the corneal edema caused by the trauma gradually resolved over time (Fig. 4B and 4C), with these conditions persisting until the end of the study.

By the first month post-surgery, the wound sizes in all four groups decreased over time. Utilizing the advantage of fluorescein sodium directly adhering to damaged corneal surfaces, a series of non-invasive examinations and visual analyses were performed to quantify the corneal wounds in different groups. The 8AB group (0.071 mm²) and 10AB group (0.001 mm²) showed nearly complete corneal wound healing after 2 months recovering, while the control and suture groups remained respectively at 1.132 mm² and 3.647 mm² (Fig. 4E and 4F). The wound area in the suture group was larger than that in the blank control group, indicating that although mechanical stabilization via sutures facilitates temporary wound closure, the persistent interfacial abrasion between sutures and corneal epithelium induces microtrauma, thereby elevating the propensity for localized inflammatory cascades and pathological angiogenesis. Compared to the control and suture groups, the corneas treated with Mfp hydrogels exhibited greater transparency and less neovascularization during the healing process (Fig. 4D).

Anterior segment optical coherence tomography (AS-OCT) images further revealed that the hydrogels tightly connected the wound edges, completely filling the defect (Fig. 5A). Compared to the control and suture groups, the hydrogel-treated corneas exhibited curvature closer to normal, reducing the likelihood of irregular astigmatism caused by improper suturing^[26]^. Additionally, AS-OCT scans of corneal cross-sections validated the preliminary efficacy of hydrogel wound adhesion. Measurements and quantification using ImageJ showed that the proportion of full-thickness corneal wound area in the hydrogel groups decreased from 22.46% (8AB) –21.84% (10AB) on day 1 post-surgery to 5.35% (8AB) –5.28% (10AB) at 2 months post-surgery (Fig. 5B). Since corneal perforating injuries are severe, keratocytes around the wound inevitably differentiate into opaque myofibroblasts^[27]^. Although fibrosis scars were almost invisible under slit-lamp examination and fluorescein sodium staining, AS-OCT detected subtle fibrotic changes in the corneal cross-sections (Fig. 5A). In the control group, the formation of adherent leukoma caused iris adhesion to the corneal inner surface, resulting in high-reflectivity masses in AS-OCT images. In contrast, the suture, 8AB, and 10AB groups exhibited weaker and smaller high-reflectivity masses (Fig. 5A and 5B).

**Fig. 5.**
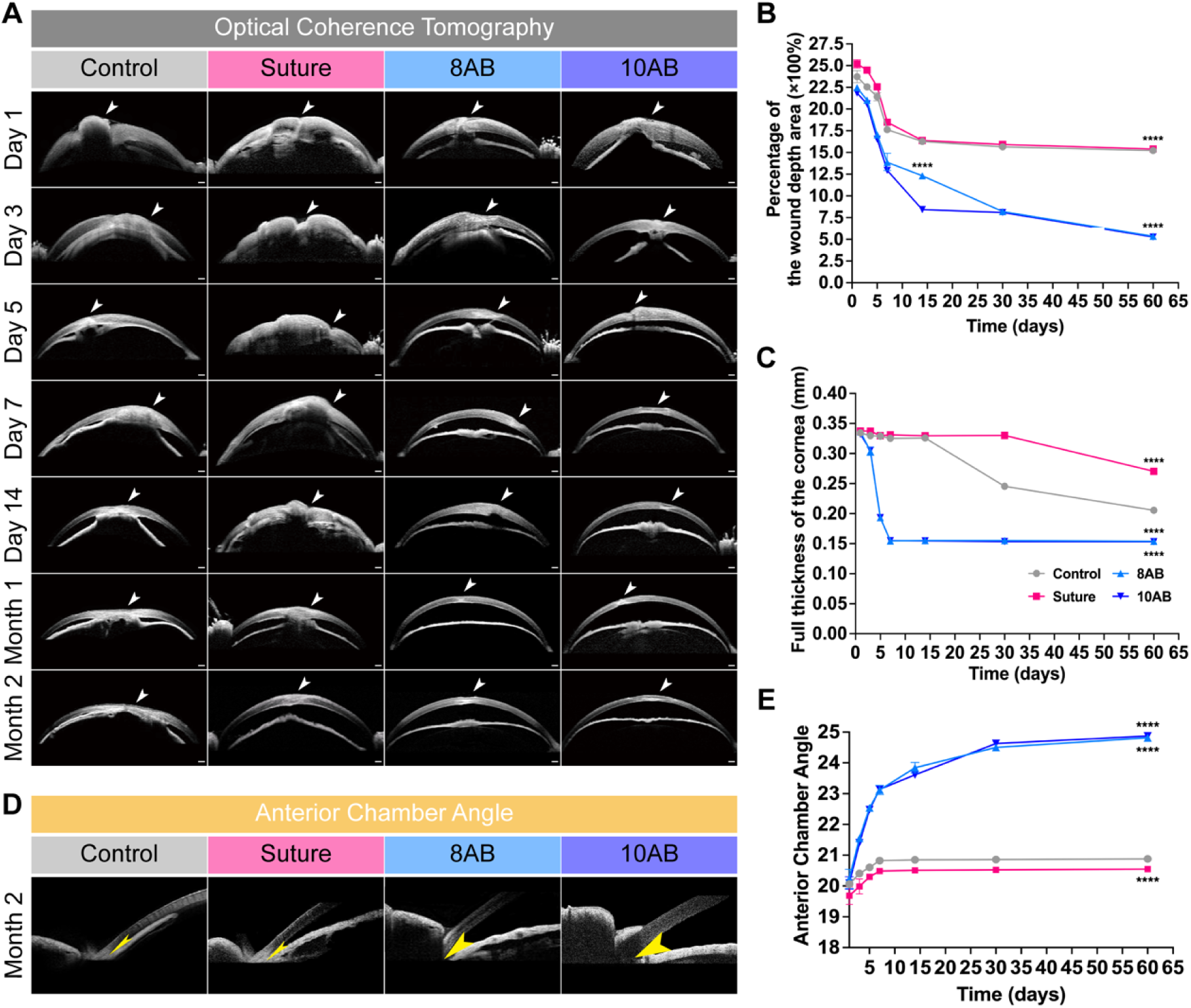
Longitudinal assessment of corneal healing via anterior segment OCT (AS-OCT) in a rat corneal perforation model. **(A)** Representative AS-OCT images showing key healing parameters at multiple timepoints (1 day to 2 months post-surgery): wound closure (white arrows), corneal-iris apposition, surface regularity, and stromal fibrosis. Scale bar: 200 µm. **(B)** Sagittal wound area expressed as percentage of total corneal thickness. **(C)** Quantitative analysis of corneal edema resolution and thickness normalization. **(D)** Comparative AS-OCT images at 2 months postsurgery demonstrating failed anterior chamber restoration with iris prolapse and adherent leukoma in the control group and preserved anterior chamber anatomy (yellow arrows indicate angles) in the treatment groups (suture, 8AB, 10AB). Scale bar: 200 µm. **(E)** Anterior chamber angle measurements using AS-OCT software (n=5). Statistical analysis by two-way ANOVA; \*\*\*\**P*<0.0001 versus control.

Additionally, the efficacy of hydrogel wound closure was demonstrated by changes in anterior chamber angles observed in AS-OCT images. After perforating injury, the anterior chamber angles in all groups decreased (> 20%) (Fig. 4B). Although the control group reflected the natural healing process post-corneal trauma, the eyes remained hypotonic, and the anterior chamber failed to normalize (Fig. 5D). Since anterior chamber recovery depends on the physiological production of aqueous humor, immediate improvement in anterior chamber angles was not expected. The hydrogel groups showed significant increases in anterior chamber angles approximately 3–5 days post-surgery, with statistically significant improvements compared to the control group within 1 month of hydrogel placement (*P* < 0.0001, Fig. 5E). The mechanical performance of wound closure in different hydrogel concentrations was statistically consistent (*P* > 0.05, *P* = 0.3997).

### Long-Term Biocompatibility of Mfp Hydrogels in Corneal Wound Repair

Since the purpose of this experiment is to investigate the long-term safety and biocompatibility of hydrogels *in vivo*, rats at the 6-month post-surgery mark were selected to detect potential long-term adverse effects. Tissue cross-sections with the perforation and hydrogel placement sites marked (Fig. 6A, white arrows) show the wound site and hydrogel placement location. After 6 months, the corneas had partially restored their original structure. The control group exhibited iris adhesion, consistent with previous AS-OCT scan results (Fig. 5A and 6A). In the suture group, due to the sutures not being removed, typical acute inflammatory reactions formed a porous fibrous layer around the suture sites, resulting in a thicker cornea compared to the other three groups (Fig. 5C). In the 8AB group, due to the inferior mechanical properties of the hydrogel compared to 10AB, the edges of the corneal perforation remained visible. Trichrome staining revealed that the original hydrogel site was covered by a mixed layer of new fibrous tissue and collagen, with no signs of iris adhesion (Fig. 6A, black arrows). Additionally, the 10AB group demonstrated nearly complete corneal regeneration due to its excellent mechanical properties, with trichrome staining confirming a clear corneal structure devoid of inflammatory cells or fibrous layers.

**Fig. 6.**
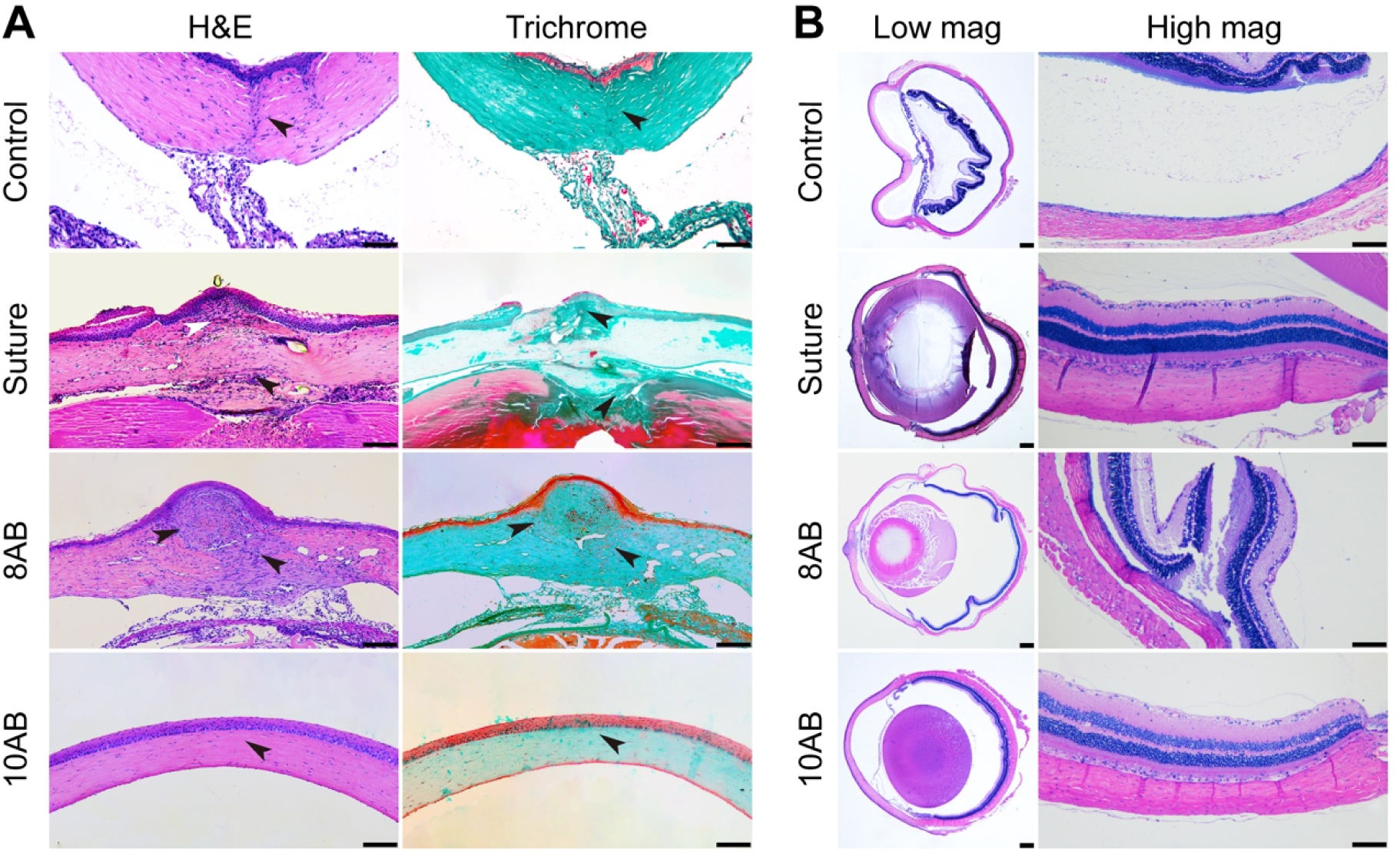
Histological evaluation of corneal tissue reorganization 2 months post-injury. **(A)** Representative H&E and Masson’s trichrome-stained corneal cross-sections from different treatment groups. Scale bar: 50 µm. **(B)** Retinal histology showing low-magnification overview of retinal architecture (*left panel*) and high-magnification details confirming the absence of trauma-induced choroidal/retinal detachment or neurotoxicity (*right panel*). Scale bar: 50 µm.

In all four groups, there was no evidence of retinal or choroidal detachment after 6 months, nor any signs of optic nerve toxicity (Fig. 6B, *left*). Hematoxylin and eosin (H&E) staining evaluation showed that the retinal layers in the experimental and suture groups exhibited no degeneration of photoreceptor outer segments or disorganization of other layers. The ganglion cell layer, inner and outer nuclear layers, and choroid with blood vessels were consistent with those in the control group. The anterior segment structures were more clearly visible than in the control group (Fig. 6B, *right*). Based on these series of cellular and animal experiments, 10AB was selected as the candidate material for subsequent *in vivo* experiments in New Zealand white rabbits.

### Adhesion and Wound Healing in Perforating Corneal Trauma of New Zealand Rabbits

New Zealand white rabbits were divided into two groups for the experiment, with the suture group serving as the control. Due to the thinness of rat corneas, AS-OCT could not accurately discern the true state of the hydrogel within the cornea. New Zealand white rabbits addressed this limitation, as AS-OCT scans of corneal cross-sections within 1-day post-surgery showed the hydrogel successfully fixed between corneal layers and connecting the wound edges (Fig. S2), providing a foundation for rapid healing. At 14 days post-surgery, AS-OCT further revealed that the wound in the 10AB group was smaller and fully integrated with the corneal defect, with only the elastic layer of the cornea curling inward due to trauma (Fig. 7A). Under normal light, neither group exhibited corneal neovascularization caused by trauma or foreign bodies. Slit-lamp examination showed recovery of the anterior chamber in both the suture and 10AB groups. Additionally, fluorescein sodium staining indicated that the suture group caused more corneal epithelial damage than the hydrogel group (Fig. 7B).

**Fig.7.**
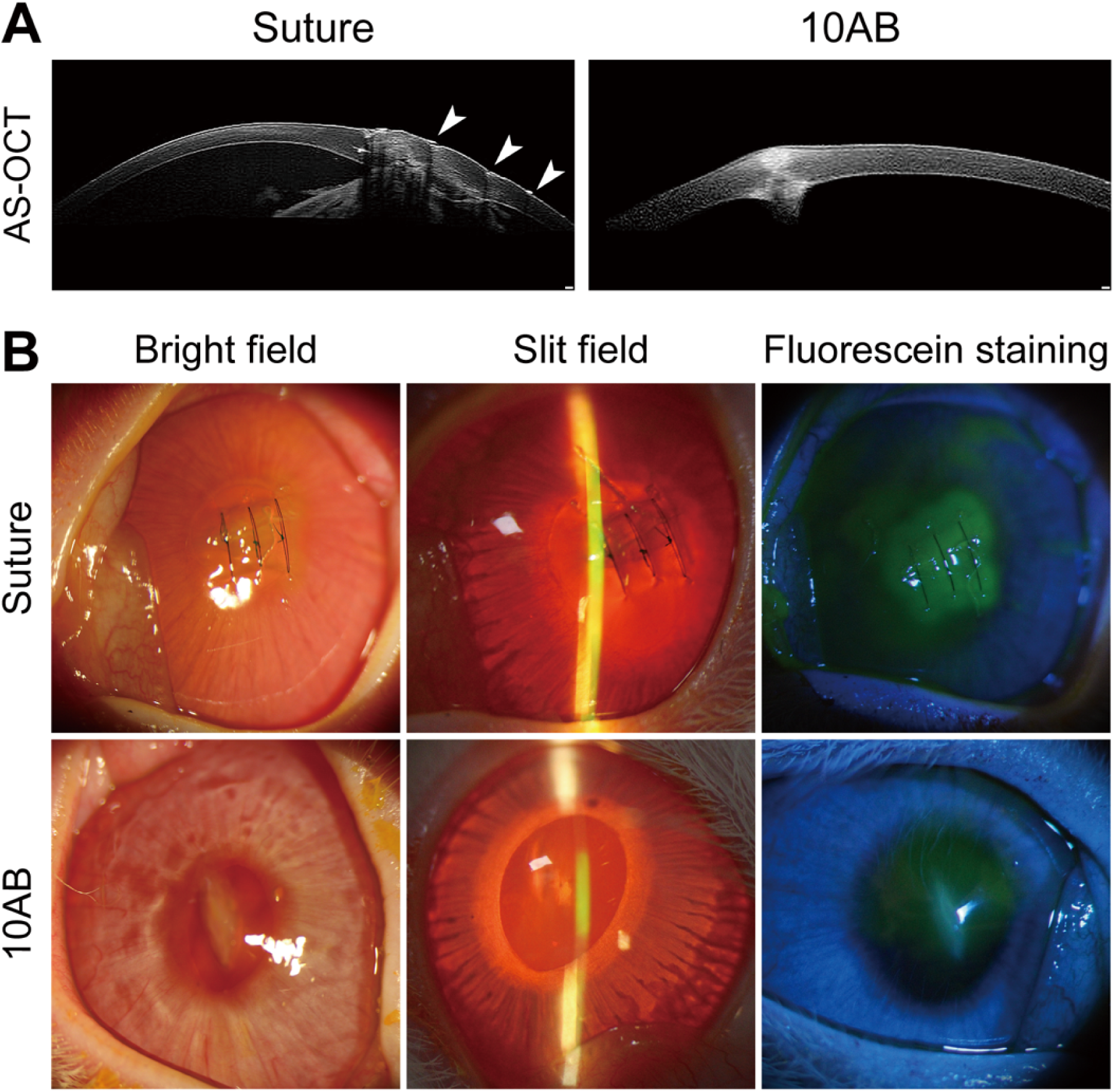
Mfp hydrogels promote wound healing in a rabbit corneal perforation model. **(A)** Comparative corneal repair at 14 days in the suture (needle tracks marked by white arrows) and 10AB hydrogel groups. Scale bar: 200 μm. **(B)** Multimodal imaging (bright field, slit-lamp microscopy, and fluorescein staining under cobalt blue light) at 14 days postsurgery.

Although our study is the first to propose a fully protein-based hydrogel for repairing perforating corneal trauma and has preliminarily demonstrated its safety and efficacy, several limitations warrant consideration. First, while the hydrogel showed promising results in rats, the testing period in larger animals like rabbits is insufficient, and the intraocular pressure in larger animals is generally higher than in small animals like rats. Therefore, long-term studies in rabbits are needed. Additionally, while the unmodified hydrogel demonstrated foundational safety and waterproof adhesion *in vivo*, this investigation did not exploit its modular protein engineering potential. Prior work established this system’s capacity for post-gelation functionalization via spherical protein conjugation and stem cell encapsulation—features critical for advanced regenerative applications^[14]^. To address this gap, we plan to encapsulate Fidgetin-Like 2 (FL-2) or LSCs within the 10AB hydrogel after repairing perforating corneal trauma in New Zealand white rabbits to validate its potential for inhibiting corneal neovascularization and promoting corneal stromal cell regeneration.

## Discussion

Mussel foot protein 3 (Mfp3), along with related materials like Mfp5 and engineered 3,4-dihydroxyphenyl-L-alanine (DOPA)-containing proteins^[28]^, exhibit a unique combination of properties: robust interfacial adhesion, intrinsic photothermal activity, biocompatibility, and adaptive mechanical behavior under physiological conditions. ^[18]^. These characteristics make Mfp-based protein materials highly promising for biomedical applications, including tissue engineering, surgical adhesives, and drug delivery^[29,30]^.Building upon these attributes, our group previously engineered the recombinant fusion hydrogel SpyCatcher-ELP-Mfp3-ELP-SpyCatcher (BMB)^[14]^. This architecture features an Mfp3 core flanked by elastin-like polypeptide (ELP) domains and terminal SpyCatcher modules^[14,15]^. The ELP domains confer enhanced solubility, mechanical resilience, and structural flexibility, while the SpyCatcher modules enable site-specific covalent crosslinking via SpyTag interactions. Our previous studies reported that this recombinant protein improves upon conventional Mfp hydrogels, achieving 93% human mesenchymal stem cell (hMSC) viability over 7 days and strong adhesion to biological substrates like porcine skin^[14]^.

In this study, our group introduced a recombinant protein construct expressed in E. coli: 6×His-SpyTag-ELP-SpyTag-ELP-SpyTag (AAA)^[15]^. AAA and BMB were enzymatically crosslinked via Streptomyces antibioticus tyrosinase at defined molar ratios to generate tunable hydrogels. Comparative analysis revealed that the 10 wt% formulation (10AB) exhibited superior mechanical properties, including enhanced stiffness (elastic modulus: 10AB > 8AB), stronger interfacial adhesion (24.3 ± 4.7 vs. 21.4 ± 6.0 N/cm²), and improved degradation resistance (<15% mass loss over 7 days in vitro), attributable to its denser crosslinked network relative to 8AB.

In cellular experiments, both hydrogels maintained LSC stem cell activity, and 10AB exhibited more cellular forces and adhesion sites than 8AB in vertical cell migrations. While 8AB allowing more cell migration in through its porous network due to its lower crosslinking density. Therefore, both hydrogel formulations, each exhibiting distinct advantages, were employed to assess their therapeutic efficacy in animal models. In animal models, 10AB demonstrated superior waterproof adhesion and repair performance under the complex physicochemical conditions of the cornea, where the outer surface is exposed to air and the inner surface is bathed in aqueous humor. Compared to the suture group, 10AB showed lower rates of inflammation caused by foreign bodies and better restoration of corneal curvature. Our findings demonstrated that 10AB achieved effective sealing of perforating corneal trauma via its intrinsic cross-linking capability, independent of external light-curing or temperature modulation, while simultaneously preserving retinal structural integrity and visual function.

In summary, these findings underscore the translational potential of Mfp hydrogels. Unlike suture-based approaches, which rely heavily on surgical precision, the intrinsic material adaptability of Mfp hydrogel—including its programmable gelation kinetics and self-adhesive properties—mitigates surgeon-dependent variability while offering error-tolerant application^[31]^. Beyond demonstrating efficacy in acute corneal repair, this platform shows significant therapeutic potential for addressing trauma-induced neovascularization, offering dual regenerative and anti-pathological capabilities.

## Materials and Methods

### Materials

The pQE80l and pACYCDuet plasmids was purchased from Qiagen and Addgene, respectively. The creation of the gene constructs, *SpyTag-ELP-SpyTag-ELP-SpyTag* (*AAA*) and *SpyCatcher-ELP-MFP-3-ELP-SpyCatcher* (*BMB*) has been described previously^[14,15]^. The porcine skin was purchased from a local supermarket (ParknShop). Chemicals and regents were purchased from Sangon Biotech.

### Production and purification of recombinant proteins

*E. coli* strain BL21 (DE3) was used for protein expression. To produce the recombinant proteins, the bacterial cells harboring the corresponding plasmids, pQE80l::*SpyTag-ELP-SpyTag-ELP-SpyTag* and pQE80l::*SpyCatcher-ELP-MFP-3-ELP-SpyCatcher* were grown in LB at 37°C till the mid-log phase (the optical density at 600 nm or OD_600_ is ∼0.6 to 0.8), followed by the addition of 1 mM isopropyl-β-d-thiogalactopyranoside (IPTG; Sangon Biotech) to induce protein expression at 37°C. After 2 hours, the cells were harvested and flash-frozen in liquid nitrogen. The proteins were purified using Ni– nitrilotriacetic acid chromatography following the manufacturer’s recommendations (GE Healthcare Inc.). The purified proteins were dialyzed against Milli-Q water (5 liters × 4) at 4°C and lyophilized at −80°C. The resulting protein powders were stored at −80°C before use.

Tyrosinase from *Streptomyces antibioticus* was cloned, expressed, and purified using standard recombinant protein expression methods^[14,32]^. The tyrosinase gene was purchased as gBlocks from Integrated DNA Technologies, cloned into pACYCDuet plasmid, and expressed in E. coli BL21(DE3) in TB medium [yeast extract (24 g/L), tryptone (20 g/L), glycerol (4 mL/L), and 0.17 M KH_2_PO_4_ + 0.72 M K_2_HPO_4_ (100 mL/L)]. The E. coli cells harboring pACYCDuet-tyrosinase were grown to an OD_600_ of 0.6 to 0.8 and induced with 0.5 mM IPTG at 16°C for 20 hours. The cells were then harvested through centrifugation and resuspended in lysis buffer [300 mM NaCl and 50 mM tris-HCl (pH 8.0)]. The cell suspension was lysed through sonication (Bandelin SONOPULS HD 4400, UW 400, TS113). Proteins were then purified using a 5-ml HisTrap high performance (HP) column (Cytiva Life Sciences) according to the manufacturer’s recommendations. Purified tyrosinase was further dialyzed against the dialysis buffer [50 mM sodium phosphate and 0.01 mM CuSO_4_ (pH 6.5)] at 4°C, diluted to a concentration of 1 mg/mL, flash-frozen with liquid nitrogen, and then stored at −80°C before use.

### Hydrogel formation

Lyophilized AAA and BMB protein were dissolved in sterilized phosphate-buffered saline (PBS, pH 7.4) to make an 8 wt% or 10 wt % solution. The AAA and BMB protein solutions were mixed at 1:4 ratio, followed by mixing with 1 μg of tyrosinase per 30 μL mixture in an Eppendorf tube to initiate gelation at room temperature.

### Dynamic shear rheology

Rheological measurements were performed with 50 μl of hydrogels on a TA Instruments ARES-G2 strain-controlled rheometer with a customized steel parallel plate geometry (8 mm in diameter for the upper fixture and 25 mm in diameter for the bottom fixture) at room temperature. Test modes included dynamic time sweep, strain sweep and frequency sweep. Time sweep tests were performed at a fixed frequency of 5 rad/s and strain at 1%. Strain sweep tests were performed over a range of 0.01 to 1000% strain at a fixed frequency of 5 rad/s. Frequency sweep tests were performed from 100 to 0.01 rad/s by holding the strain at 1%.

### Erosion-rate measurements

Erosion of the hydrogels was assessed in DPBS at pH 7.4 under 37°C^[33]^. The gels were immersed in 1 mL of DPBS. Aliquots of 20 μL were taken at defined time points and fresh DPBS (20 μL) was added back to keep the volume constant. Protein concentrations were determined by the absorbance at 280 nm using a NANODROP 2000C spectrophotometer (Thermo Scientific).

### Shear adhesion tests

Adhesion tests on porcine skin were performed on the same rheometer. The porcine skin strips (30 mm by 10 mm by 2 mm) were obtained by cropping a piece of cleaned porcine skin. Hydrogels (10 μL) were applied on the surface (10 by 10 mm2) ofporcine skin strips (30 by 10 mm2) and then placed at 100% relative humidity at room temperature for 3 hours. This prolonged incubation under the humid condition was intended to examine the water resistance of the adhesive ELMs. Tensile tests were performed at a crosshead speed of 0.1 mm/min, and the peak forces were recorded.

### Rabbit LSC Isolation and Culture

According to our previously improved cell extraction method^[34]^, rabbit LSCs were isolated. Tissue approximately 16 mm in diameter, including the cornea, limbus, and conjunctiva, was excised from fresh New Zealand white rabbit eyeballs using a scalpel and incubated at 4°C in 2.5 mg/mL dispase solution for 12 hours. Single cells were further digested with 0.25% trypsin/EDTA for 5 minutes. After inhibition with trypsin inhibitor, the cells were centrifuged at 1000 rpm for 5 minutes to collect the cell pellet. The cells were then resuspended in LSC medium, which consisted of Defined Keratinocyte-SFM basal medium (Gibco), Defined Keratinocyte-SFM growth supplement (Gibco), and 1% antibiotic-antimycotic (Gibco).

### Characterization of Cytotoxicity

The cytotoxicity of the hydrogel formed by crosslinking BMB and AAA was evaluated based on a previous report^[14]^. Briefly, the mixed solution of BMB and AAA was spread onto the bottom of a 24-well plate. After gelation, the hydrogel was washed three times with PBS and then soaked in culture medium for 48 hours. Limbal stem cells were seeded at a density of 100 × 10 cells per well. Medium containing Defined Keratinocyte-SFM basal medium, Defined Keratinocyte-SFM growth supplement, and 1% antibiotic-antimycotic served as the positive control. Cell proliferation was first measured using the Cell Counting Kit-8 (CCK-8, Beyotime). LSCs were incubated in CCK-8 solution for 2 hours at 37°C in a 5% CO incubator after 24, 48, or 120 hours of culture. Absorbance was measured at 450 nm, with each test group containing three independent experimental samples. Cell numbers were correlated with optical density (OD). Cell viability (%) was determined using the formula: (Experimental group - Blank group) / (Control group - Blank group) × 100%. Cytotoxicity was assessed using the Calcein-AM/PI double staining kit (CA/PI, Beyotime)^[35]^. LSCs were treated appropriately for different durations (1, 3, 5, and 7 days), washed twice with PBS (Gibco), and incubated in the dark at 37°C for 30 minutes with 5 µM Calcein-AM and 10 µM PI (provided in the Calcein-AM/PI double staining kit). Stained cells were observed under a fluorescence microscope (Leica, DMi8)

### Transwell migration assay

8AB (8 wt% solution) and 10AB (10 wt% solution) were spread onto the bottom of the experimental wells without touching the upper chambers. LSCs were then placed in the upper chambers of Transwell plates (8 µm pore size, Corning, NY, USA) without growth supplement. Defined K-SFM containing 0.2% growth supplement was added to the lower chambers. Cells were treated at 37°C with 5% CO for 48 hours. The upper surfaces of the Transwell chambers were washed, and cells were fixed with 4% paraformaldehyde (Servicebio) for 30 minutes. Cells were stained with 0.1% crystal violet (Servicebio) for 20 minutes, and migrated cells in six random fields were counted under an inverted microscope.

### Rhodamine-Phalloidin Fluorescence Labeling Method

After incubating LSCs cultured in 3D on-top mode for 24 hours, the old Defined K-SFM medium was removed, and the cell-covered slides (NEST) were washed with 1 mL PBS. Cells were fixed with 4% paraformaldehyde for 10 minutes, permeabilized with 0.1% Triton X-100 (Servicebio) for 15 minutes, and then stained with rhodamine-phalloidin (YEASEN) at room temperature in the dark for 40 minutes^[36]^. After washing with PBS to remove unbound fluorescent dye, the slides were inverted onto glass slides and observed under a fluorescence microscope.

### Perforating Corneal Trauma Model

Surgical and animal care procedures were conducted according to protocols approved by the Institutional Animal Care and Use Committee of Shanghai Sixth People’s Hospital Affiliated to Shanghai Jiao Tong University School of Medicine and adhered to the guidelines of the National Institutes of Health (NIH) for animal care and use. All SD rats and New Zealand white rabbits were obtained from the animal laboratory of Shanghai Sixth People’s Hospital Affiliated to Shanghai Jiao Tong University School of Medicine. The corneal perforating injury model was adapted from Antonio Guirao et al.^[37]^. Briefly, preoperative mydriasis was induced using tropicamide eye drops to prevent iris incarceration in the corneal defect area during surgery. Rats were anesthetized by intraperitoneal injection of 60 mg/kg sodium pentobarbital solution, while rabbits were anesthetized by intravenous injection of 30 mg/kg sodium pentobarbital solution via the ear margin. Surface anesthesia was achieved using lidocaine eye drops. An infant eyelid speculum was placed to fully expose the cornea. Sodium hyaluronate viscoelastic, a colorless transparent gel solution composed of sodium hyaluronate and physiologically balanced salt solution, was injected into the anterior chamber using a 1 mL syringe to protect the iris and lens from mechanical damage. A full-thickness corneal incision was made at the central cornea using a 15° ophthalmic knife, and the incision was enlarged to approximately 3 ± 0.5 mm (rats) or 6 ± 0.5 mm (rabbits) using microforceps. Leakage of viscoelastic from the wound was considered a successful model of corneal perforating injury. Animals were randomly divided into control, suture, 8AB, and 10AB groups. The control group received wound irrigation with 0.9% saline, while the suture (10-0 suture, Ethicon, Johnson & Johnson Medical), 8AB, and 10AB groups had their wounds closed using sutures or hydrogels. The eyelids of each animal were closed for 10 minutes to allow the hydrogel to fully solidify. Postoperatively, levofloxacin eye drops were administered four times daily for 7 days to prevent infection and foreign body rejection.

### Slit-Lamp and AS-OCT Evaluation

Animals were monitored daily, and corneal wound healing was assessed on specified days (1, 3, 5, 7, 14 days, and 1 and 2 months) using a slit lamp (Zeiss) and anterior segment optical coherence tomography (AS-OCT, SPECTRALIS^®^ HRA + OCT, Heidelberg Engineering, Germany). Weekly slit-lamp examinations were performed to assess corneal opacity and neovascularization. Fluorescein sodium ophthalmic strips were used under cobalt blue light to monitor and photograph corneal epithelial staining, evaluating the extent of corneal epithelial regeneration. AS-OCT was used to obtain cross-sectional images to measure corneal thickness, longitudinal wound healing, and anterior chamber angle recovery.

### Histological Analysis

SD rats were humanely euthanized 6 months post-surgery. Enucleated eyeballs were fixed in Davidson’s fixative (Solarbio)^[38]^ to maintain structural integrity. Tissues were dehydrated using a dehydrator (Leica, HistoCore PEAR) following a preset program: 75% ethanol → 85% ethanol → 95% ethanol → absolute ethanol → xylene → paraffin. Tissues were then embedded in paraffin using an embedding machine (Leica) and sectioned. Sections were stained with hematoxylin and eosin (H&E) or Masson’s trichrome stain to evaluate local inflammation and corneal fibrosis. Choroidal or retinal detachment and the arrangement of retinal photoreceptor segments and optic nerve layers were confirmed by H&E staining and evaluated by a professional pathologist.

### Statistical Analysis

All data are presented as mean ± SD. *In vitro* experiments were performed with 3–6 technical replicates across 3 independent experiments. *In vivo* experiments included at least 6 rats per group. Statistical analysis was performed using Analysis of Variance (ANOVA), with a *P*-value less than 0.05 considered statistically significant.

## Acknowledgments

We thank Biosciences Central Research Facility of HKUST (Clear Water Bay) to support the work in general equipments and the technical assistance provided by the core facility platform at the Shanghai Sixth People’s Hospital Affiliated to Shanghai Jiao Tong University School of Medicine.

## Author contributions

Conceptualization: FS, DL, ZBZ

Methodology: QKY, LX, YLY, FS

Experiments: LX, QKY, PH, YTH, YHY, HC, HKFF

Visualization: PZ, XW

Supervision: DL, FS, ZBZ, YLY, SZK

Writing—original draft: LX, QKY

Writing—review & editing: DL, FS, ZBZ, YLY

## Data and materials availability

All data are available in the main text or the supplementary materials.”

## References

1. R. N. Palchesko, S. D. Carrasquilla, A. W. Feinberg, Natural Biomaterials for Corneal Tissue Engineering, Repair, and Regeneration. Adv Healthc Mater 7, e1701434 (2018).

2. S. Takahashi, T. Ono, K. Abe, Y. Mori, R. Nejima, T. Iwasaki, T. Miyai, K. Miyata, Prognosis and etiology of traumatic and non-traumatic corneal perforations in a tertiary referral hospital: a 30-year retrospective study. Graefes Arch Clin Exp Ophthalmol 260, 629– 635 (2022).

3. H. Chen, J. Han, X. Zhang, X. Jin, Clinical Analysis of Adult Severe Open-Globe Injuries in Central China. Front Med (Lausanne) 8, 755158 (2021).

4. M. Soleimani, K. Cheraqpour, F. Salari, K. Fadakar, S. Habeel, S. M. Baharnoori, S. Banz, S. A. Tabatabaei, F. A. Woreta, A. R. Djalilian, All about traumatic cataracts: narrative review. J Cataract Refract Surg 50, 760–766 (2024).

5. T. J. Patterson, A. Kedzierski, D. McKinney, J. Ritson, C. McLean, W. Gu, M. Colyer, S. F. McClellan, S. C. Miller, G. A. Justin, A. K. Hoskin, K. Cavuoto, J. Leong, A. Rousselot Ascarza, F. A. Woreta, K. E. Miller, M. C. Caldwell, W. G. Gensheimer, T. Williamson, F. Dhawahir-Scala, P. Shah, A. Coombes, G. Sundar, R. A. Mazzoli, M. Woodcock, S. L. Watson, F. Kuhn, S. Halliday, R. S. M. Gomes, R. Agrawal, R. J. Blanch, The Risk of Sympathetic Ophthalmia Associated with Open-Globe Injury Management Strategies: A Meta-analysis. Ophthalmology 131, 557–567 (2024).

6. J. Pelletier, A. Koyfman, B. Long, High risk and low prevalence diseases: Open globe injury. Am J Emerg Med 64, 113–120 (2023).

7. B. Gurnani, K. Kaur, “Scleral and Limbic Lacerations” in StatPearls (StatPearls Publishing, Treasure Island (FL), 2024; http://www.ncbi.nlm.nih.gov/books/NBK580563/).

8. J. Du, G.-Y. Zheng, C.-L. Wen, X.-F. Zhang, Y. Zhu, Long-term outcomes of wedge resection at the limbus for high irregular corneal astigmatism after repaired corneal laceration. Int J Ophthalmol 9, 843–847 (2016).

9. J. Dalma-Weiszhausz, M. Galván-Chávez, E. B. Guinto-Arcos, D. Y. Miyake-Martínez, A. Rodríguez-Reyes, M. F. Golzarri, C. Sebastián-Arellano, N. M. Dávila-Ávila, C. E. Ríos-Elizondo, Full-versus partial-thickness sutures: experimental models of corneal injury repair. Int Ophthalmol 41, 325–334 (2021).

10. A. Sharma, R. Kaur, S. Kumar, P. Gupta, S. Pandav, B. Patnaik, A. Gupta, Fibrin glue versus N-butyl-2-cyanoacrylate in corneal perforations. Ophthalmology 110, 291–298 (2003).

11. J. Stark, M. de Leval, Experience with fibrin seal (Tisseel) in operations for congenital heart defects. Ann Thorac Surg 38, 411–413 (1984).

12. B. Duchesne, H. Tahi, A. Galand, Use of human fibrin glue and amniotic membrane transplant in corneal perforation. Cornea 20, 230–232 (2001).

13. S. Masket, J. A. Hovanesian, J. Levenson, F. Tyson, W. Flynn, M. Endl, P. A. Majmudar, S. Modi, R. Chu, M. B. Raizman, S. S. Lane, T. Kim, Hydrogel sealant versus sutures to prevent fluid egress after cataract surgery. J Cataract Refract Surg 40, 2057–2066 (2014).

14. Q. Lu, E. Danner, J. H. Waite, J. N. Israelachvili, H. Zeng, D. S. Hwang, Adhesion of mussel foot proteins to different substrate surfaces. J R Soc Interface 10, 20120759 (2013).

15. J. Luo, X. Liu, Z. Yang, F. Sun, Synthesis of Entirely Protein-Based Hydrogels by Enzymatic Oxidation Enabling Water-Resistant Bioadhesion and Stem Cell Encapsulation. ACS Appl Bio Mater 1, 1735–1740 (2018).

16. F. Sun, W.-B. Zhang, A. Mahdavi, F. H. Arnold, D. A. Tirrell, Synthesis of bioactive protein hydrogels by genetically encoded SpyTag-SpyCatcher chemistry. Proc Natl Acad Sci U S A 111, 11269–11274 (2014).

17. H. J. Kang, F. Coulibaly, F. Clow, T. Proft, E. N. Baker, Stabilizing isopeptide bonds revealed in gram-positive bacterial pilus structure. Science 318, 1625–1628 (2007).

18. L. Li, J. O. Fierer, T. A. Rapoport, M. Howarth, Structural analysis and optimization of the covalent association between SpyCatcher and a peptide Tag. J Mol Biol 426, 309–317 (2014).

19. A. V. Ljubimov, M. Saghizadeh, Progress in corneal wound healing. Prog. Retin. Eye Res. 49, 17–45 (2015).

20. K. L. Xu, N. Di Caprio, H. Fallahi, M. Dehghany, M. D. Davidson, L. Laforest, B. C. H. Cheung, Y. Zhang, M. Wu, V. Shenoy, L. Han, R. L. Mauck, J. A. Burdick, Microinterfaces in biopolymer-based bicontinuous hydrogels guide rapid 3D cell migration. Nat. Commun. 15, 2766 (2024).

21. T. Yeung, P. C. Georges, L. A. Flanagan, B. Marg, M. Ortiz, M. Funaki, N. Zahir, W. Ming, V. Weaver, P. A. Janmey, Effects of substrate stiffness on cell morphology, cytoskeletal structure, and adhesion. Cell Motil. Cytoskelet. 60, 24–34 (2005).

22. M. Ehrbar, A. Sala, P. Lienemann, A. Ranga, K. Mosiewicz, A. Bittermann, S. C. Rizzi, F. E. Weber, M. P. Lutolf, Elucidating the role of matrix stiffness in 3D cell migration and remodeling. Biophys. J. 100, 284–293 (2011).

23. Y. Wang, G. Wang, X. Luo, J. Qiu, C. Tang, Substrate stiffness regulates the proliferation, migration, and differentiation of epidermal cells. Burns 38, 414–420 (2012).

24. C. G. Priya, T. Prasad, N. V. Prajna, V. Muthukkaruppan, Identification of human corneal epithelial stem cells on the basis of high ABCG2 expression combined with a large N/C ratio. Microsc Res Tech 76, 242–248 (2013).

25. G. Su, X. Guo, L. Xu, B. Jin, Y. Tan, X. Zhou, W. Wang, X. Li, S. Wang, G. Li, Isolation and characterization of rabbit limbal niche cells. Exp Eye Res 241, 109838 (2024).

26. S. Feizi, M. A. Javadi, F. Javadi, P. Malekifar, H. Esfandiari, Suture-related complications after deep anterior lamellar keratoplasty for keratoconus. Graefes Arch Clin Exp Ophthalmol 262, 1195–1202 (2024).

27. R. R. Mohan, D. Kempuraj, S. D’Souza, A. Ghosh, Corneal stromal repair and regeneration. Prog Retin Eye Res 91, 101090 (2022).

28. Z. Yang, F. Sun, Self-Assembly and Genetically Engineered Hydrogels. Adv. Biochem. Eng. Biotechnol. 178, 169–196 (2021).

29. W.-Y. Quan, Z. Hu, H.-Z. Liu, Q.-Q. Ouyang, D.-Y. Zhang, S.-D. Li, P.-W. Li, Z.-M. Yang, Mussel-Inspired Catechol-Functionalized Hydrogels and Their Medical Applications. Molecules 24, 2586 (2019).

30. M. Cui, S. Ren, S. Wei, C. Sun, C. Zhong, Natural and bio-inspired underwater adhesives: Current progress and new perspectives. APL Mater. 5, 116102 (2017).

31. W. Zhang, S. Liu, L. Wang, B. Li, M. Xie, Y. Deng, J. Zhang, H. Zeng, L. Qiu, L. Huang, T. Gou, X. Cen, J. Tang, J. Wang, Triple-crosslinked double-network alginate/dextran/dendrimer hydrogel with tunable mechanical and adhesive properties: A potential candidate for sutureless keratoplasty. Carbohydr Polym 344, 122538 (2024).

32. Q. Yi, X. Dai, B. M. Park, J. Gu, J. Luo, R. Wang, C. Yu, S. Kou, J. Huang, R. Lakerveld, F. Sun, Directed assembly of genetically engineered eukaryotic cells into living functional materials via ultrahigh-affinity protein interactions. Sci Adv 8, eade0073 (2022).

33. Z. Yang, Y. Yang, M. Wang, T. Wang, H. K. F. Fok, B. Jiang, W. Xiao, S. Kou, Y. Guo, Y. Yan, X. Deng, W.-B. Zhang, F. Sun, Dynamically Tunable, Macroscopic Molecular Networks Enabled by Cellular Synthesis of 4-Arm Star-like Proteins. Matter 2, 233–249 (2020).

34. Z. Zhou, D. Long, C.-C. Hsu, H. Liu, L. Chen, B. Slavin, H. Lin, X. Li, J. Tang, S. Yiu, S. Tuffaha, H.-Q. Mao, Nanofiber-reinforced decellularized amniotic membrane improves limbal stem cell transplantation in a rabbit model of corneal epithelial defect. Acta Biomater 97, 310–320 (2019).

35. Y. Yu, H. Xiao, G. Tang, H. Wang, J. Shen, Y. Sun, S. Wang, W. Kong, Y. Chai, X. Liu, X. Wang, G. Wen, Biomimetic hydrogel derived from decellularized dermal matrix facilitates skin wounds healing. Mater Today Bio 21, 100725 (2023).

36. J. A. Cooper, Effects of cytochalasin and phalloidin on actin. J Cell Biol 105, 1473–1478 (1987).

37. A. Guirao, Theoretical elastic response of the cornea to refractive surgery: risk factors for keratectasia. J Refract Surg 21, 176–185 (2005).

38. N. Bayat, Y. Zhang, P. Falabella, R. Menefee, J. J. Whalen, M. S. Humayun, M. E. Thompson, A reversible thermoresponsive sealant for temporary closure of ocular trauma. Sci Transl Med 9, eaan3879 (2017).

