## Supplemental Figures and Table for "Genetically Programmable Adhesive Protein Hydrogels for Sealing Perforating Corneal Trauma"

Ling Xu; Qikun Yi; Ping Hu *et al.*

**This PDF file includes:**

Figs. S1 to S2

Tables S1

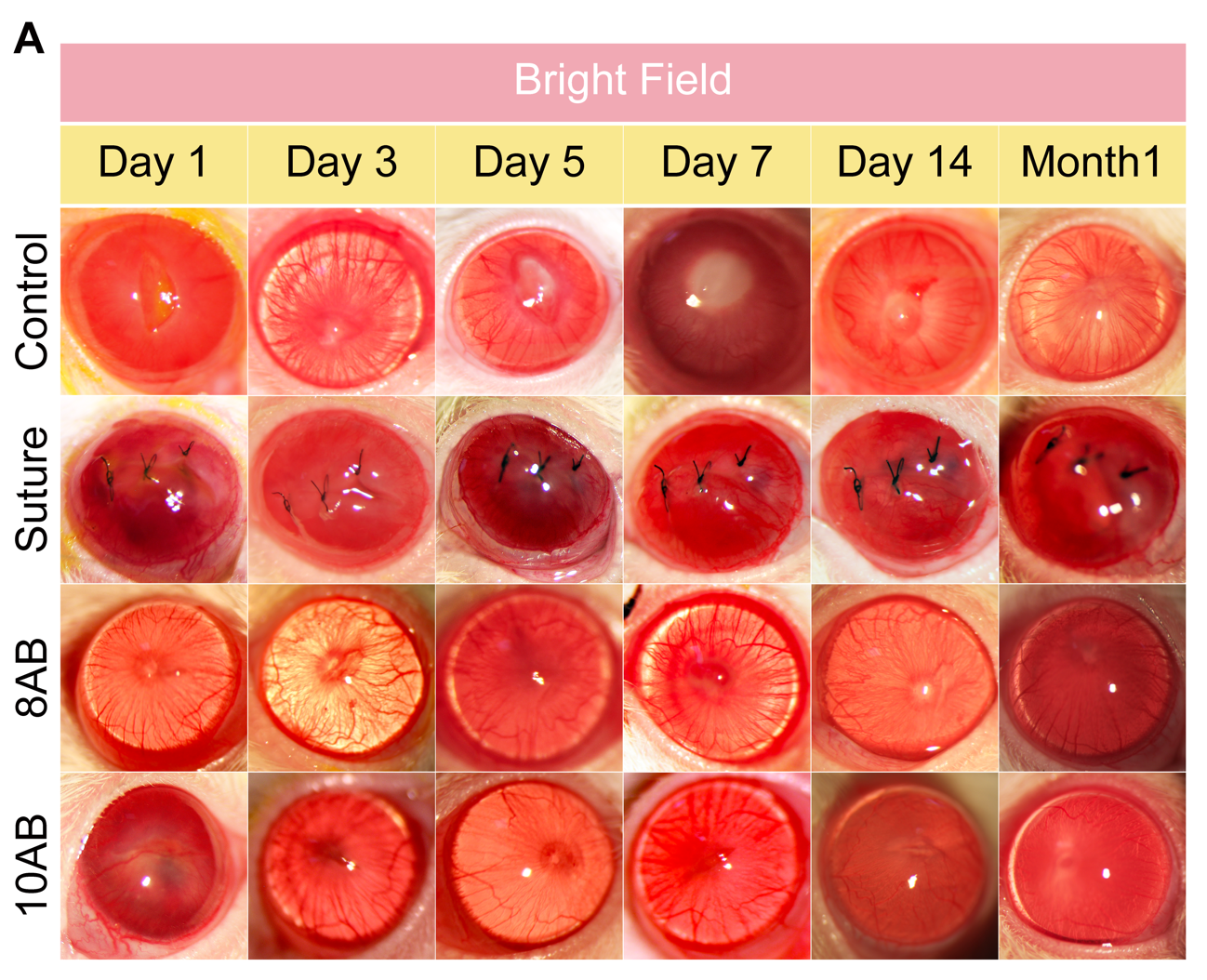

Fig. S1.

**Mfp hydrogels promote wound healing in a rat corneal perforation injury model. (A)** Serial anterior segment ophthalmoscope under bright field showing wound healing in control, suture, 8AB, and 10AB groups during the 4-week follow-up.

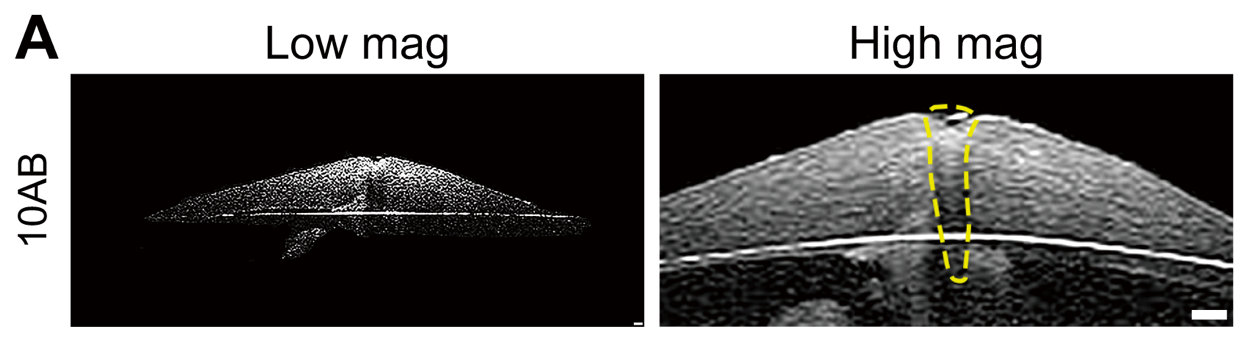

Fig. S2.

**Mfp hydrogels promote wound healing in a rabbit corneal perforation model.** **(A)** AS-OCT images at 24 hours post-surgery showing global corneal morphology (*left*) and precise hydrogel localization (yellow dashed lines outline implant; *right*). Scale bar: 200 μm.

| Protein | Amino acid sequence |
| --- | --- |
| SpyCatcher-ELP-Mfp3-ELP-SpyCatcher | MKGSSHHHHHHVDIPTTENLYFQGAMVDTLSGLSSEQGQSGDMTIEEDSATHIKFSKRDEDGKELAGATMELRDSSGKTISTWISDGQVKDFYLYPGKYTFVETAAPDGYEVATAITFTVNEQGQVTVNGKATKGDAHIDGPQGIWGQLEGHGVGVPGVGVPGVGVPGEGVPGVGVPGVGVPGVGVPGVGVPGEGVPGVGVPGVGVPGVGVPGVGVPGEGVPGVGVPGVGELMELADYYGPKYGPPRRYGGGNYNRYGRRYGGYKGWNNGWKRGRWGRKYYGRGDSDGPQGIWGQGTGTSGGSMTSVPGVGVPGVGVPGEGVPGVGVPGVGVPGVGVPGVGVPGEGVPGVGVPGVGVPGVGVPGVGVPGEGVPGVGVPGVGVPGGLVDIPTTENLYFQGAMVDTLSGLSSEQGQSGDMTIEEDSATHIKFSKRDEDGKELAGATMELRDSSGKTISTWISDGQVKDFYLYPGKYTFVETAAPDGYEVATAITFTVNEQGQVTVNGKATKGDAHIDGPQGIWGQLEWKK |
| SpyTag-ELP-SpyTag-ELP-SpyTag | MKGSSHHHHHHVDAHIVMVDAYKPTKLDGHGVGVPGVGVPGVGVPGEGVPGVGVPGVGVPGVGVPGVGVPGEGVPGVGVPGVGVPGVGVPGVGVPGEGVPGVGVPGVGELAHIVMVDAYKPTKTSVPGVGVPGVGVPGEGVPGVGVPGVGVPGVGVPGVGVPGEGVPGVGVPGVGVPGVGVPGVGVPGEGVPGVGVPGVGVPG GLLDAHIVMVDAYKPTKLEWKK |
| Tyrosinase | MGSSHHHHHHSQDMTVRKNQASLTAEEKRRFVAALLELKRTGRYDAFVTTHNAFILGDTDNGERTGHRSPSFLPWHRRFLLEFERALQSVDASVALPYWDWSADRSTRSSLWAPDFLGGTGRSRDGQVMDGPFAASAGNWPINVRVDGRTFLRRALGAGVSELPTRAEVDSVLAMATYDMAPWNSGSDGFRNHLEGWRGVNLHNRVHVWVGGQMATGVSPNDPVFWLHHAYIDKLWAEWQRRHPSSPYLPGGGTPNVVDLNETMKPWN DTTPAALLDHTRHYTFDVVDH |

Table S1.

**Amino acid sequences of recombinant proteins used in this study.**
